# Evolution-guided engineering of an ancient nitrogenase interface enhances enzyme activity and stability

**DOI:** 10.64898/2026.06.05.730521

**Authors:** Elias I. Kemna, Rajdeep Banerjee, Betül Kaçar

**Affiliations:** Department of Bacteriology, University of Wisconsin – Madison, Madison, WI

## Abstract

Nitrogenase is the only enzyme capable of biological nitrogen fixation and a major target for sustainable agricultural engineering, yet its functional and structural complexity has made it difficult to identify regions that can be modified without compromising activity or assembly. Here, we use structural evolution to identify a lineage-specific N-terminal extension in NifK as a candidate engineering target within the MoFe protein interface. By generating variant libraries exceeding 9,000 members and evaluating them through deep mutational scanning, diazotrophic growth assays and in vitro characterization to map the sequence-function landscape of this interface across variable conditions. We show that NifK extension is required for nitrogenase activity while remaining broadly tolerant to mutation, revealing a sequence-function landscape that is both flexible and constrained. A subset of residues at the NifD-NifK interface are critical for maintaining complex stability. Structural analyses indicate that the extension stabilizes the MoFe complex via co-evolved electrostatic interactions, and targeted mutations can improve both enzymatic activity and thermostability. These findings identify the NifK extension as a tunable interface, providing a strategy for engineering more robust nitrogenase variants.

## Introduction

Biological nitrogen fixation emerged early in Earth’s history, with geochemical and phylogenetic evidence placing its origin approximately 3 billion years ago (Stüeken et al., 2015; Garcia et al., 2020). This process is mediated by nitrogenase, which remains the only known enzymatic mechanism capable of reducing atmospheric nitrogen to ammonia (Zhang et al., 2020). Prior to the development of Haber–Bosch process, biological nitrogen fixation supplied virtually all new reactive nitrogen to agricultural systems (Fowler et al., 2013), underscoring the central role of nitrogenase in both global food production and environment (Zhang et al., 2020). Current efforts to introduce nitrogenase into eukaryotic hosts offer a potential route toward sustainable nitrogen fertilizer production but remain hindered by challenges including host incompatibility and oxygen sensitivity (Dixon & Kahn, 2004; Bennett et al., 2023). Accordingly, nitrogenase represents a prime target for protein engineering.

Nitrogenase catalysis is coordinated by two metalloproteins: the electron-donor Fe protein, and the catalytic MoFe protein. The Fe protein is a homodimer containing a [4Fe-4S] cluster, whereas the MoFe protein is a heterotetramer containing two catalytic metalloclusters. The M-cluster (FeMo-cofactor; [MoFe₇S₉C-homocitrate]) serves as the active site for dinitrogen reduction, while the P-cluster ([Fe₈S₇]) mediates electron transfer from the Fe protein to the M-cluster during catalysis (Einsle and Rees, 2020). Understanding how nitrogenase structure influences these complex interactions and relates to function is therefore a central challenge in nitrogenase engineering.

The exploration of sequence-function landscapes is a cornerstone of protein engineering. However, these landscapes are highly dimensional and rugged (Katznacheev, 2019), and the combinatorial space of possible variants exceeds experimental capacity, making it difficult to prioritize mutations or regions of interest (Fowler & Fields, 2014). This challenge is amplified in multi-protein complexes such as nitrogenase, where function depends on coordinated interactions between subunits (Guntas et al., 2010; Taggart and Li, 2019; Hsia et al., 2021). Evolutionarily informed approaches that integrate phylogenetic and structural information provide a powerful framework for identifying regions of functional importance and restricting exploration to information-rich regions.

Our recent survey of nitrogenase structural evolution (Cuevas-Zuviria et al., 2025) identified several lineage-specific motifs specific to certain classes of nitrogenase. Among these, structural phylogenetics demonstrated an extended NifK N-terminus, almost exclusively associated with Nif-I nitrogenases (**Fig. 1a**), a lineage composed predominantly of aerobic and facultative bacteria (Garcia et al., 2020). Consistent with this observation, ancestral sequence reconstruction revealed a progressive elongation of the NifK N-terminus along evolutionary trajectories leading to extant Nif-I nitrogenases (Cuevas-Zuviria et al., 2025; Garcia et al., 2023). Such stepwise elongation and long-term conservation of the NifK N-terminus strongly suggest that this feature was selectively maintained during Nif-I evolution, yet the functional consequences of its emergence and evolution remain unknown.

**Fig. 1:**
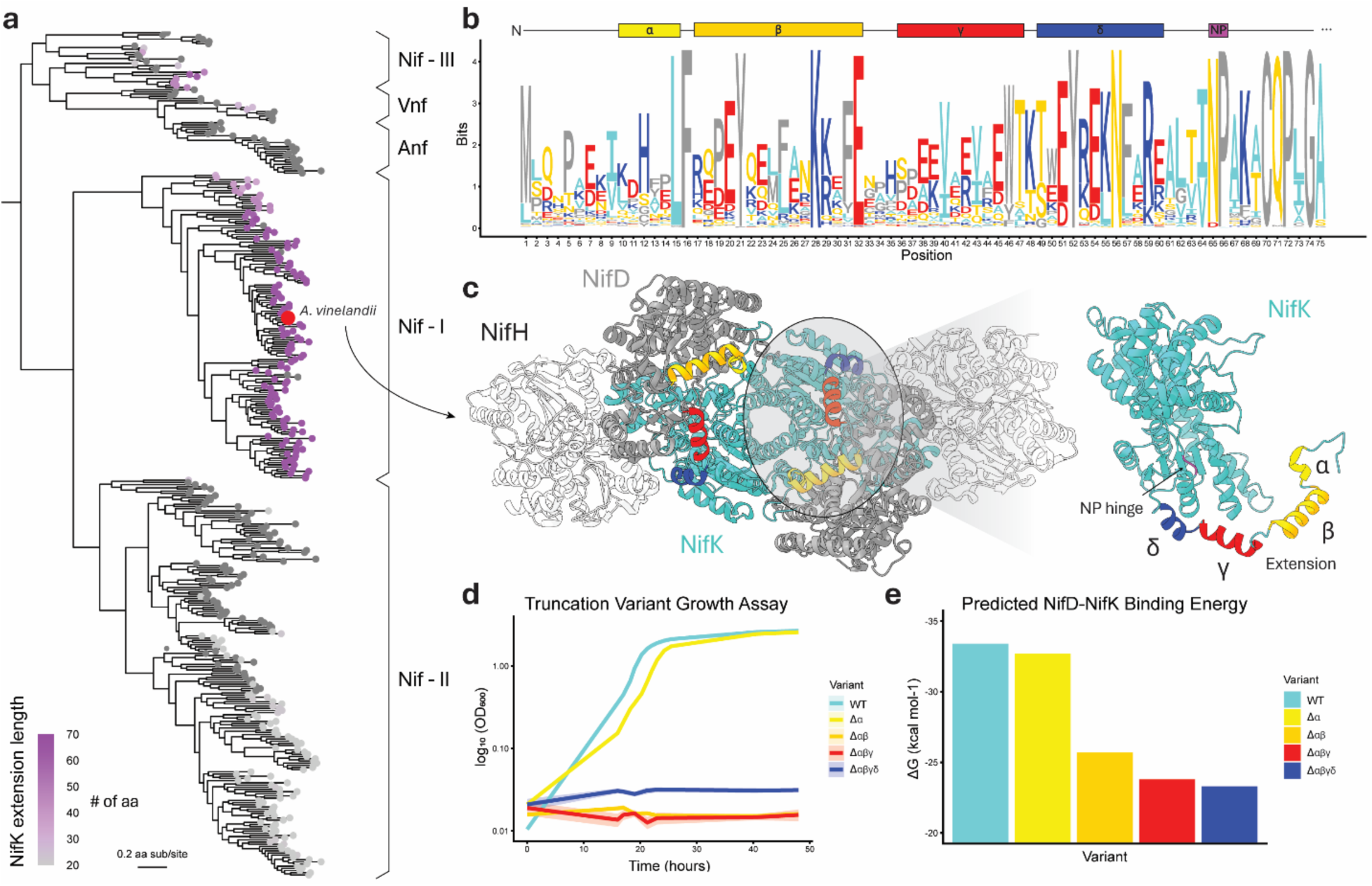
The NifK N-terminus extension is essential for nitrogenase function. **(a)** Nitrogenase phylogeny built from concatenated Nif/Vnf/AnfHDK protein sequences (Cuevas Zuviría et al., 2025). Tip labels colored by length of NifK extension (# of aa). **(b)** Sequence logo illustrating amino acid frequencies for each position from a multiple sequence alignment of 639 extant NifK N-terminus extensions, with secondary structure annotated above. **(c)** Structure of *A. vinelandii* nitrogenase complex (PDB 4WZB) colored by subunit. The NifK subunit is highlighted, with the N-terminus extension labeled. **(d)** Diazotrophic growth of NifK truncated variants. The truncated region is indicated; data represent mean ± s.d. (n=3 biological replicates). **(e)** Predicted NifD-NifK binding affinities of the NifK extension truncation variants.

In this study, we developed a high-throughput nitrogenase variant assembly and screening platform that enabled systematic exploration of the NifK sequence-function landscape through the evaluation of more than 9,000 unique variants. By combining deep mutational scanning, random mutagenesis, and targeted functional characterization, we systematically explored the sequence-function landscape of the NifK N-terminal extension in model organism *Azotobacter vinelandii* under multiple conditions. Integrating structural analyses and predictive modeling with empirical measurements, we demonstrate that the NifK extension evolved into a key component of the NifD–NifK interface, providing stabilizing interactions within the MoFe protein complex. Biochemical and biophysical characterization of representative variants further validated the mechanistic basis of these interactions.

## Results

### Structural evolution of the NifK extension

We defined the boundaries of the NifK extension based on a multiple sequence alignment of 639 extant nitrogenase sequences (**Fig. 1b**). Approximately 65 residues from the N-terminus, we observe a highly conserved asparagine-proline motif (referred to as the NP hinge hereafter) that introduces a structural bend and turns the protein structure away from the core of NifK (**Fig. 1c**). In contrast, residues before the NP hinge are less conserved. Considering this, we define the NP hinge as the boundary of the NifK N-terminus extension. The *Azotobacter vinelandii (A. vinelandii*) N-terminus NifK extension spans 66 residues in length and consists of four α-helical segments, designated the Alpha, Beta, Gamma and Delta helices (**Fig. 1c**).

To probe the function of the NifK N-terminus extension, we systematically created various truncations of the *A. vinelandii* extension by shortening the extension one helix at a time and then assessed these variants for growth in N_2_-fixing conditions (**Fig. 1d**). Removal of the Alpha helix results in a slight growth defect, extending the lag phase by ∼3 hours, which progresses to complete loss of growth upon subsequent deletion of the Beta helix. This loss of growth is not recovered by subsequent deletion of the Gamma or Delta helices. Additionally, predicted binding energies for the helix truncation variants reveal a slight decrease in binding energy with loss of the Alpha helix (2%), followed by a more dramatic decrease with loss of the Beta helix (23%), coinciding with loss of growth (**Fig. 1e**).

### Evolution of a NifD – NifK electrostatic interface

Building upon the observed correlations between extension length and NifD-NifK binding energies, we closely analyzed the NifD-NifK extension interface. Interfacing residues between NifD and NifK in the *A. vinelandii* MoFe nitrogenase structure (PDB: 3U7Q) were predicted using PDBePISA (see Supplementary Table). Of the 66 amino acids comprising the extension, 41 are predicted to interface with NifD; 15 forming hydrogen bonds and 4 forming salt bridges. These results indicate that the extension contributes a significant portion of the NifD–NifK interface. It accounts for 25.4% (15/59) of predicted hydrogen bonds, 20% (4/20) of predicted salt bridges, and 35.3% (41/116) of total interfacing residues, despite representing only 12.6% (66/524) of NifK.

To observe how the interacting surface of NifD changes over time, we generated electrostatic maps for ancestral NifD proteins, comparing surface charge patterns to NifK extension length (**Fig. 2a, b**). In the earliest ancestral reconstructions, the electrostatic surface of NifD differs markedly from that of extant *A. vinelandii*, with negative charge broadly distributed across the protein surface. In Anc_2 and Anc_3, coinciding with the first substantial elongation of the NifK extension, the negative charge becomes more localized, concentrating near the N-terminus of the extension. From Anc_4 through Anc_5, as the extension continues to lengthen, the negative charge further redistributes, remaining localized at the N-terminal region of the extension. From Anc_6 to extant *A. vinelandii*, the extension does not increase in length, and the pocket stays in place, building up in negative charge. This migration of negative charge relative to the N-terminus of the extension indicates a possible co-evolution of the NifD and NifK extension interface.

**Fig. 2:**
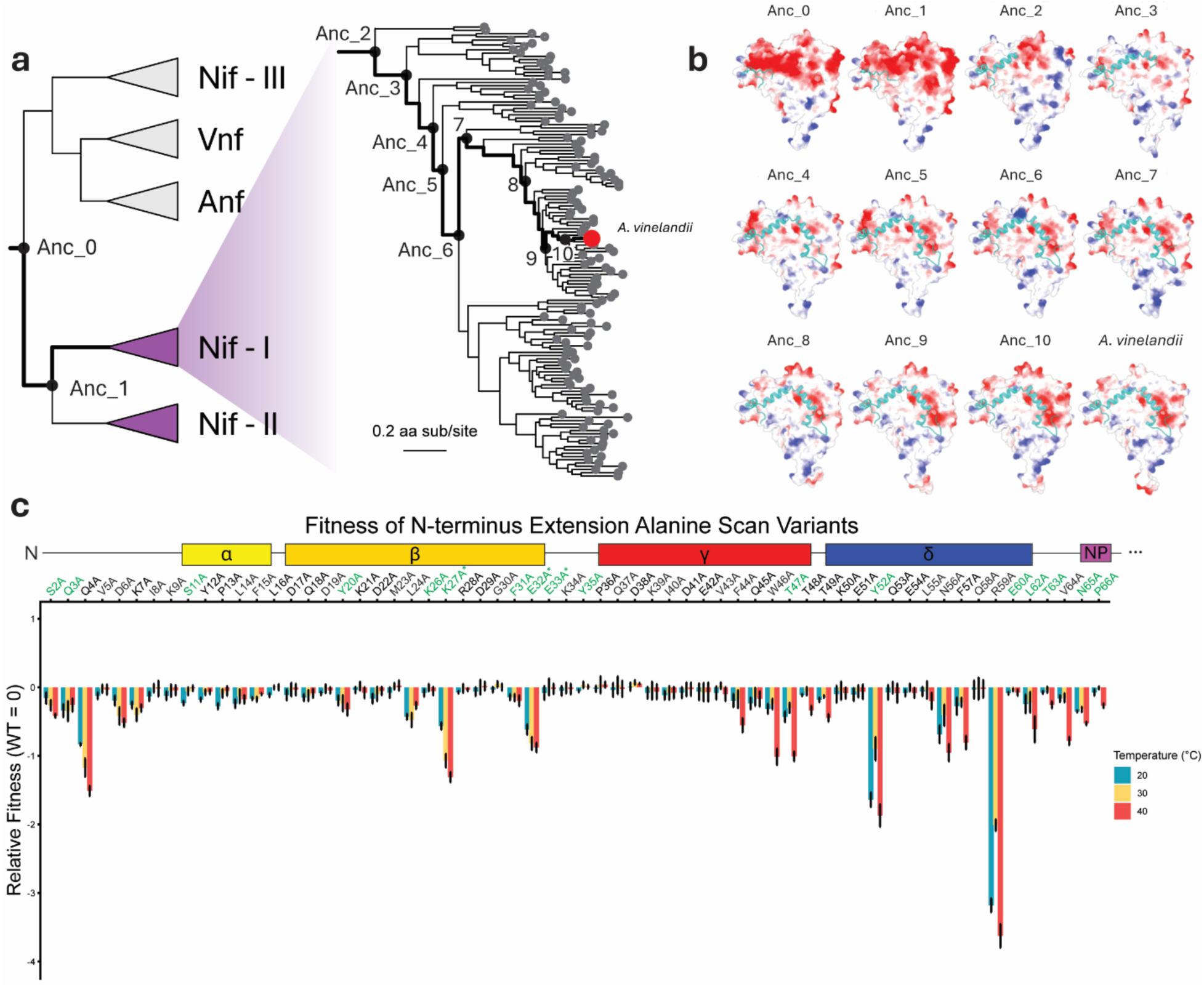
Evolution and interactions of the NifD-NifK interface. **(a)** Simplified nitrogenase phylogeny highlighting major clades with Nif-I clade in detail. Ancestral nodes leading to *A. vinelandii* are indicated as dots and labeled by Anc number. **(b)** Electrostatic surface maps of NifD for representative ancestral and extant variants, with the corresponding NifK extension colored sea-foam green. Negative and positive charges are shown in red and blue, respectively. **(c)** Alanine scan of the NifK extension. The x-axis represents individual alanine point mutants along the extension, with secondary structure annotated above. The y-axis shows relative fitness compared to wild type (WT). Values represent the mean of three biological replicates ± s.d. Residues predicted by PDBePISA to form hydrogen or ionic bonds with the interface of NifD are highlighted in green. Ionic bonds are indicated with asterisks.

### NifK extension alanine scan

To further test the co-evolution of NifD and NifK interfaces, we generated a NifK extension alanine scan library, creating single alanine substitutions for each residue in the extension, and assessed competitive diazotrophic growth at three different temperatures (20, 30, and 40 °C) to probe the mutational tolerance for each position (**Fig. 2c**). Of the 65 variants tested, 18 demonstrated significant fitness defects as compared to WT at 30 °C. Of those defective variants, 77.8% (14/18) interface with NifD (see Supplementary Table). Additionally, all defective variants, besides Y52A and R59A, perform significantly worse at 40 °C than 20 °C. These results confirm that residues interfacing with NifD are especially important for the function of the NifK extension and mutations at these residues often result in sensitivity to high temperature conditions; further indicating that their role is involved in inter-subunit stability. Additionally, we observe that for most of the residues along the extension (72.3%; 47/65) mutations to alanine do not result in a significant fitness change.

### Fitness characterization of the NifK extension

To explore the sequence-function landscape of this NifK extension and assess its potential as an engineering target, we generated a deep mutational scan library of the NifK extension and quantified competitive diazotrophic growth across three temperatures (20, 30 and 40 °C) (**Fig. 3a, b**). Of the 1,236 possible single-amino acid variants, we were able to construct and obtain enough sequencing reads to determine the relative fitness for 1,233 variants (99.8%).

**Fig. 3:**
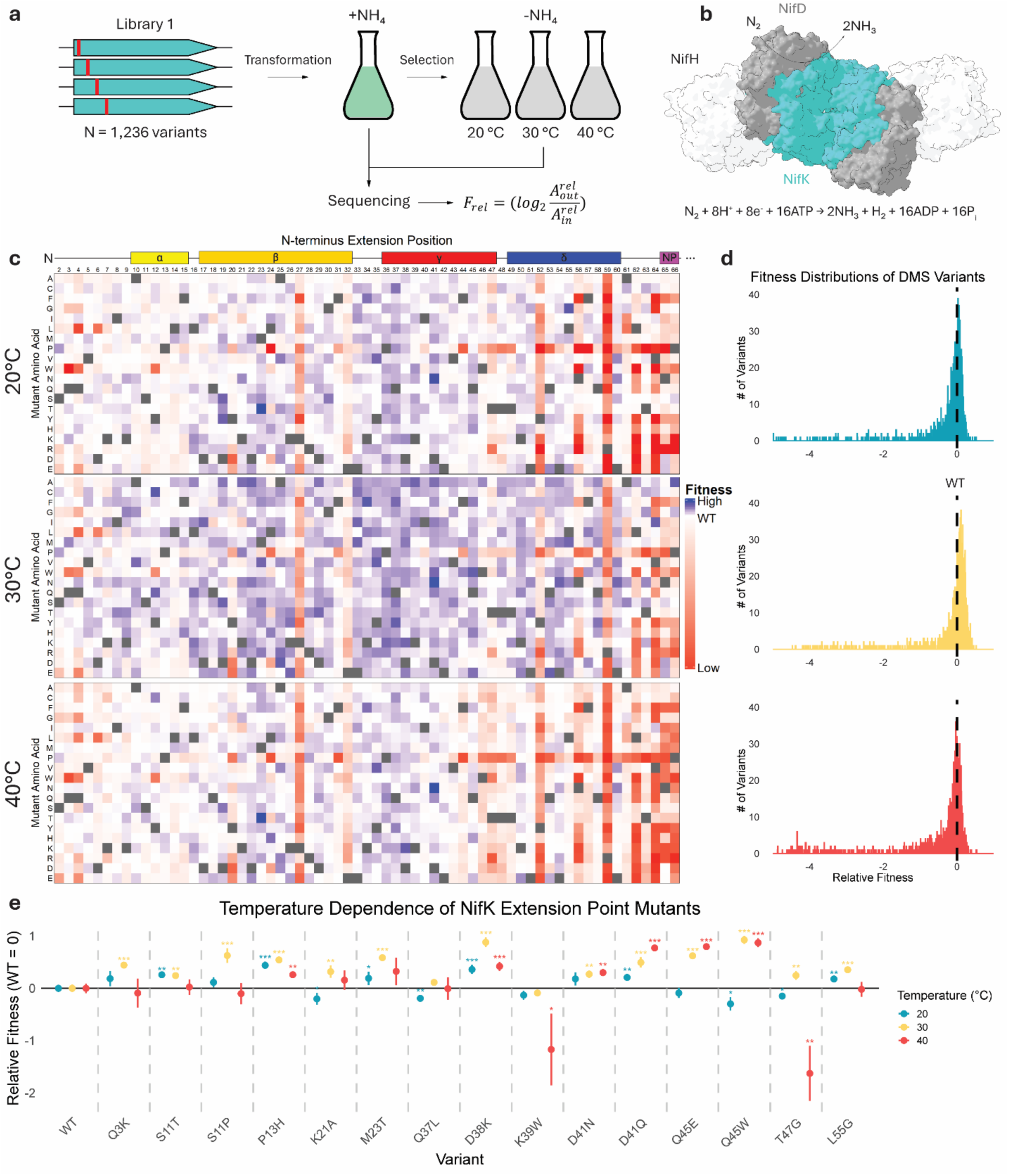
Fitness charcterization of the NifK N-terminus extension. **(a)** Schematic of deep mutational scanning, pooled diazotrophic growth assays and sequencing. **(b)** Architecture of *A. vinelandii* molybdenum nitrogenase (PDB: 4WZB) and chemical equation of dinitrogen reduction catalyzed by molybdenum nitrogenase. **(c)** Heatmaps showing the fitness scores for all single amino acid substitutions at 20, 30 and 40 °C. The x-axis indicates extension position, with secondary structure annotated above, and y-axis indicates substituted amino acid identity. Fitness values represent the mean of three biological replicates. Color scale denotes relative fitness compared to wild type (WT): blue, greater than WT; white, equal to WT; red, less than WT. Dark gray squares indicate the WT amino acid at each position. **(d)** Distribution of fitness values for all point mutants at 20, 30 and 40 °C. The black dashed line indicates WT fitness. **(e)** Fitness of individually tested point mutants realtive to wild type (WT) at 20, 30 and 40 °C. Values represent the mean of four biological replicates ± s.d. Statistical significance relative to WT is indicated by asterisks (*p < 0.05, **p < 0.01, ***p < 0.001).

Our results indicate that the NifK extension is broadly tolerant of mutation; most variants are neutral or modestly beneficial. Across all three temperatures, fitness values follow a unimodal distribution, with 30°C supporting the greatest number of beneficial variants (**Fig. 3c, d**). These findings differ from typical DMS results, which have a much higher proportion of detrimental variants (Sarkisyan et al., 2016; Prywes et al., 2024).

The most mutation-tolerable positions are located on the solvent-exposed regions of the NifK extension and do not make any observable interactions with NifD (**Fig. S1a**). However, several are observed at interfacing positions (positions 8, 23, 26, 30, 33, 34, 35, 37, 39, 60). We observe that certain residues interacting with NifD are largely intolerant to mutation, including K27, E32, Y52, N56, R59, L62, V64, N65, and P66. Interestingly, we identify regions of the extension that are particularly intolerable of mutation at higher temperatures (40 °C): F44, W46, T47, T48, T49, F57, and T63 (**Fig. 3c**). Upon closer structural analysis, the F44, W46, and F57 residues appear to be involved in aromatic-aromatic interactions with other NifK residues: F435 and H457, Y52, Y422 and H429, respectively (**Fig. S1b**).

When comparing the variant fitness scores across temperatures, we observe multiple cases in which certain point mutants affect fitness in a temperature-dependent manner (**Fig. 3c**). By comparing the variant fitness scores between temperature extremes (20 and 40 °C), we visualized each variant based on its temperature tolerance and sensitivity. We found multiple variants that are tolerant of low temperatures but sensitive to high temperatures or vice versa. For example, variant T47G has a fitness value of +0.12 at 20 °C, but a value of -0.45 at 40 °C (**Fig. S2**). To further investigate temperature-dependent effects, we selected 15 variants for individual testing that displayed improved fitness at a specific temperature but not others, as well as variants that demonstrate enhanced fitness across all tested temperatures (**Fig. 3e**).

Overall, these point variants retained temperature-dependent behavior, though not always in a manner consistent with the DMS library (**Table S1**). In total, roughly half (8/15) of the variants displayed the same temperature dependence during individual testing as they did during pooled testing. These discrepancies may arise from differences in how fitness was quantified in competitive versus individual growth assays. Specifically, fitness values from competitive growth assays reflect both growth rate and lag phase, whereas individual growth assays only reflect growth rate.

From individual growth assays, we observe substantial improvements in growth rate from several single point mutants as compared to wild type at all three temperatures. The top performers at each temperature consist of variant P13H, which grows ∼36% faster than WT at 20 °C, and variant Q45W, which grows ∼90% and 83% faster than WT at 30 °C and 40 °C, respectively (**Fig. 3e**).

Notably, Q45W introduces an additional aromatic residue within the high temperature-sensitive region of the NifK N-terminus extension (**Fig. 3c**). This substitution may promote the formation of additional aromatic–aromatic interactions within the protein, specifically between W45 and F44, and between W45 and F462 (**Fig. S1**). Such interactions could contribute to enhanced stability.

### NifK extension lengthening

Having identified point mutations within the NifK extension that confer improved growth relative to wild type, we next asked whether further fitness gains could be achieved by increasing the length of the extension. To test this, we constructed an extension library in which the NifK extension was elongated by inserting three random amino acids between positions Q3 and Q4. This was accomplished using three degenerate NNK codons encoded within a primer pair, enabling comprehensive sampling of sequence space at the insertion site. Additionally, we subjected this library to competitive diazotrophic growth across three different temperatures (20, 30, and 40 °C) to assess the fitness of each variant (**Fig. 4a**). Out of the 8,000 possible amino acid combinations, we were able to obtain enough sequencing reads for 7,512 variants (93.9%). From the lengthening library, we observe similar fitness distributions as seen with the DMS library. As before, there is a unimodal distribution for all three temperatures, most variants being neutral or slightly beneficial, with 30 °C supporting the largest number of beneficial mutations (**Fig. 4b**).

**Fig. 4:**
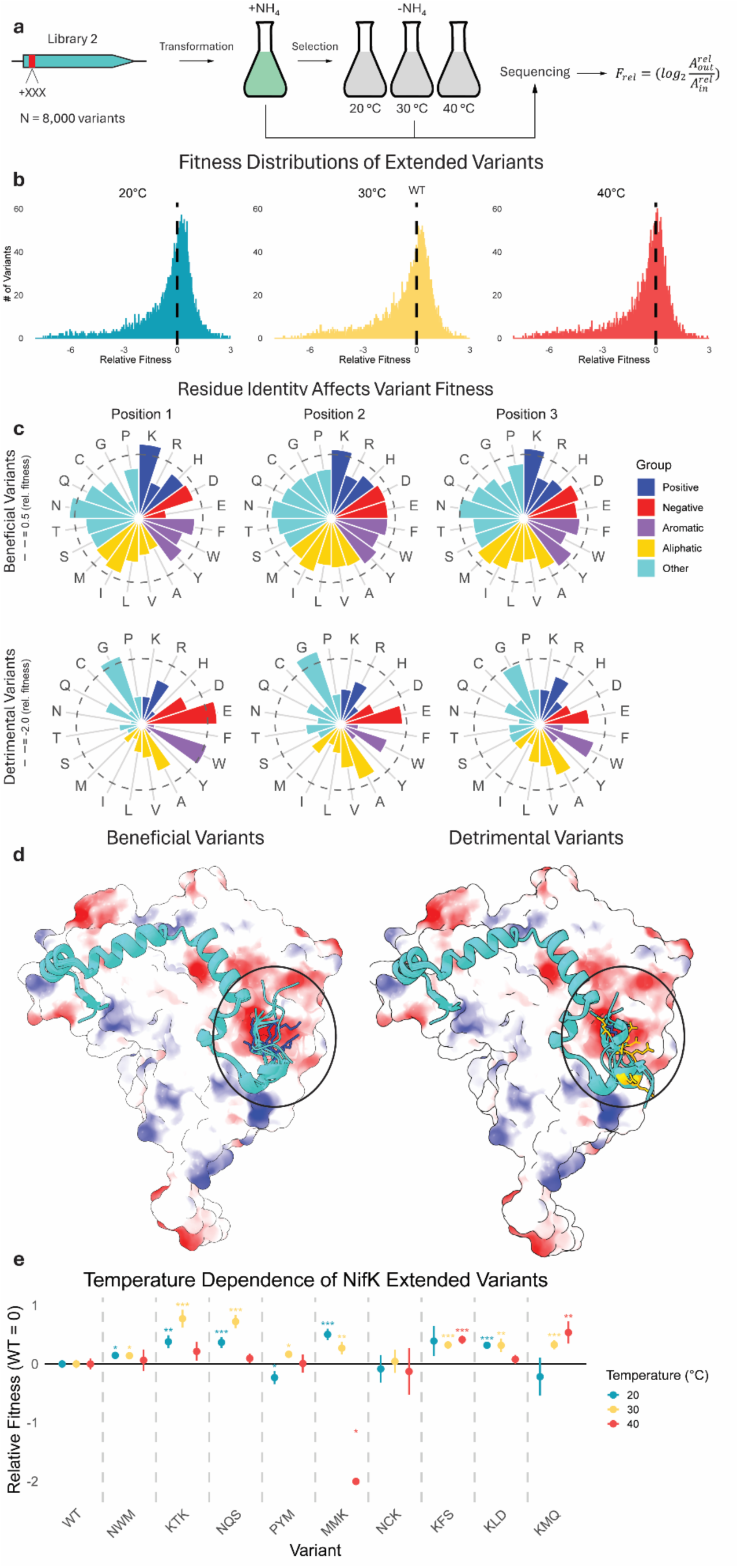
Fitness effects of NifK extension lengthening are influenced by residue identity and charge. **(a)** Schematic of triple insertion, pooled diazotrophic growth assays and sequencing. **(b)** Distribution of mean fitness values for all tested extended variants at 20, 30, and 40 °C. The black dashed line indicates wild type (WT) fitness. **(c)** Radar plots showing the average fitness contribution of each residue at each position. Variants are separated into beneficial (relative fitness > 0) and detrimental (relative fitness < 0) groups. **(d)** Predicted structures of NifK extension variants overlaid onto the electrostatic surface of NifD for the top three beneficial and detrimental variants at each temperature. Introduced lysine residues are shown as blue sticks, and glutamate residues as orange sticks. **(e)** Growth rates of individually tested lengthening mutants relative to WT at 20, 30, and 40 °C. Values represent the mean of four biological replicates ± s.d. Statistical significance relative to WT is indicated by asterisks (*p < 0.05, **p < 0.01, ***p < 0.001).

Across all variants, we found that incorporation of N, F, D, or K into the extension yields the largest fitness gains, whereas incorporation of E, G, A, or W produces the most pronounced fitness losses (**Fig 4c, Table S2**). These effects are consistent across both position and temperature for K, N, or G, while the impacts of D, F, E, A, or W are context dependent. Notably, the addition of the positively charged residue lysine uniformly enhances fitness across all positions and temperatures, whereas the negatively charged residue glutamate consistently reduces fitness. To further probe this apparent charge-dependent effect, we examined the structural localization of these three residues.

Closer examination of predicted structures shows that the added residues tend to extend toward a negatively charged pocket of NifD (**Fig. 4d**). We identified the three most significantly beneficial or detrimental variants at each temperature, defined as those with the highest or lowest fitness scores and Benjamin–Hochberg procedure-adjusted p-values < 0.1. We then superimposed these variants onto the NifD electrostatic surface and observed certain structural differences between the two groups. Beneficial variants always extend into the NifD negatively charged pocket and remain as loops while detrimental variants sometimes orient the extension away from the pocket (∼22%) and frequently form new helices (∼56%) (**Fig. 4d**). Beneficial variants are also enriched for positively charged lysine (∼67%), enabling stabilizing positive–negative electrostatic interactions, while detrimental variants often contain negatively charged glutamate (∼67%), leading to destabilizing negative–negative repulsion.

Guided by previously observed patterns, we found that nitrogenase performance can be improved by lengthening the N-terminus of NifK. To further investigate how these lengthened variants improve function, we individually tested the top three beneficial variants from the pooled NNK library at each temperature to validate the fitness estimates obtained from the pooled screen relative to WT (**Fig. 4e, Table S1**). Similarly to the previously tested point mutants, lengthened mutants often display temperature-dependent behavior. Many mutants performed significantly better than WT at one temperature but were comparable to or significantly worse than WT at another (**Fig. 4e**). For example, MMK is the top-performing at 20 °C (rel. fitness +0.50), yet the poorest performer at 40 °C (-2.00), while KMQ shows the opposite trend, performing best at 40 °C (+0.54) and worst at 20 °C (-0.22). Remarkably, lengthening the extension by just three amino acids can dramatically improve nitrogenase performance over a range of temperatures. Among the top performers, MMK grows ∼42% faster than WT at 20 °C, KTK grows ∼71% faster than WT at 30 °C, and KMQ grows ∼46% faster than WT at 40 °C (**Fig. 4e**).

### Combining beneficial NifK mutations

To assess whether beneficial point mutations and insertions have an additive effect, three double mutants were generated for each temperature. We generated new variants by either combining the two top-performing point mutations or by combining the top-performing extended mutation with one of the two best-performing point mutants. (**Fig. S3**). None of the nine double mutants exhibited an additive increase in fitness. In most cases, fitness is comparable to that of the corresponding single mutants or was slightly reduced (**Fig. S3**). These results suggest that the beneficial effects of individual mutations are not necessarily additive in this system.

### NifK extension variant activity and stability characterization

After establishing that the NifK extension plays a critical role in nitrogenase function, can be engineered to confer temperature-dependent fitness effects, and likely contributes to inter-subunit stability, we next sought to elucidate its underlying mechanism through structural and functional analyses. Four NifK extension variants were selected for purification and in vitro characterization: wild type *A. vinelandii* (WT), the Q45W point mutant, the KMQ lengthening mutant, and the Beta deletion mutant.

Given the exclusive association of the NifK extension with Nif-I nitrogenases, and the observed temperature dependencies of these variants, we sought to determine their sensitivities to oxygen and heat stress. Anaerobically purified MoFe protein complexes (**Fig. 5a**) were subjected to stress by exposing them either to atmospheric oxygen (∼21% O₂) or elevated temperature (40 °C) for varying durations, followed by measuring their enzymatic activity and oligomeric state (**Fig. 5b-e**). Remarkably, under non-stress conditions, the KMQ variant exhibits nearly twice the activity as WT MoFe (**Fig 5b, d; Fig. S4**). As expected, the Beta deletion variant shows no detectable activity under any condition. Despite initial differences, all active variants rapidly lose activity upon oxygen exposure, resulting in minimal activity within two hours. A comparable trend was observed under heat stress, with activities declining to near-equivalent levels within one hour.

**Fig. 5:**
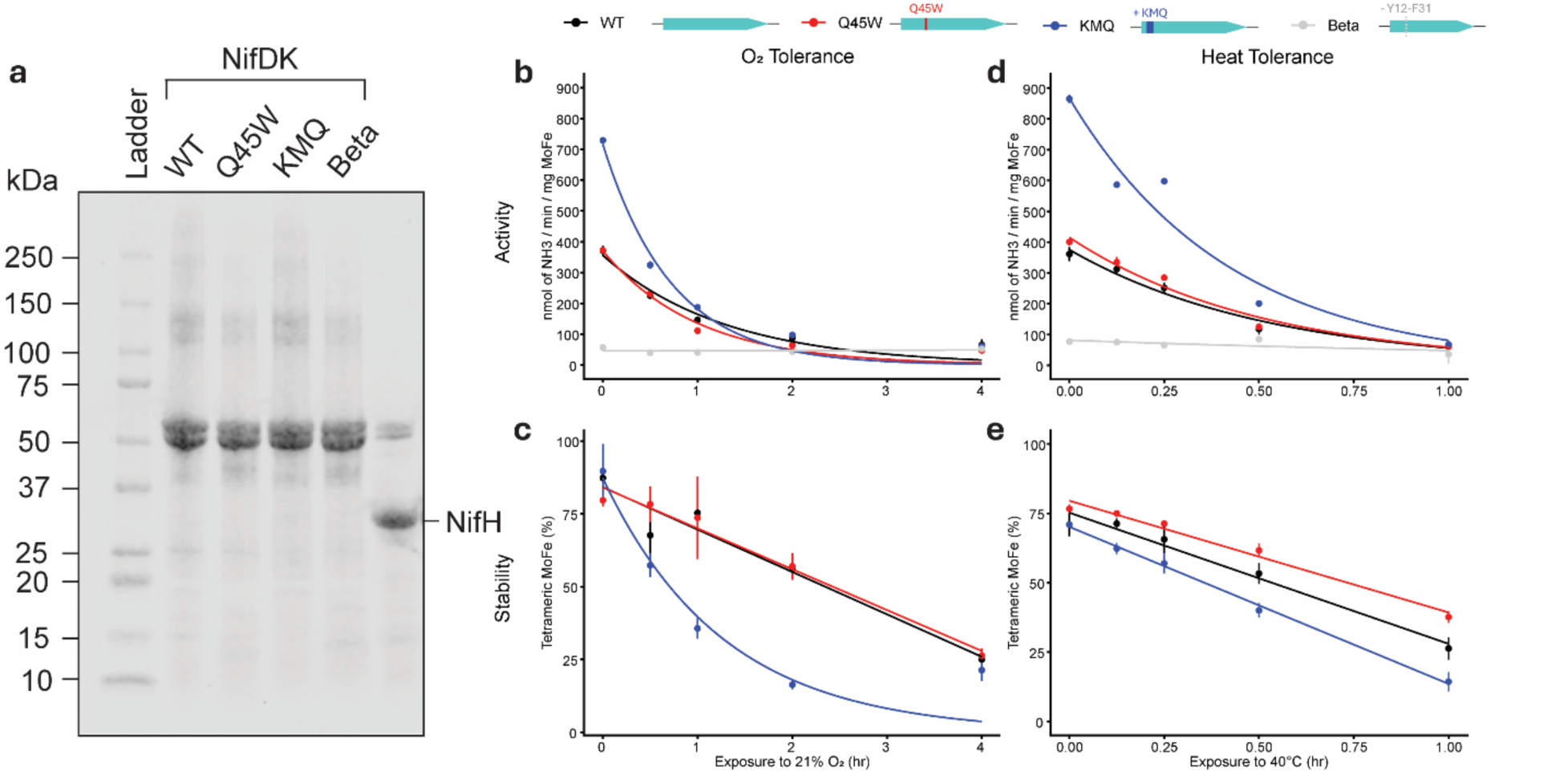
In vitro characterization of purified nitrogenase variants under stress. **(a)** SDS–PAGE analysis of anaerobically purified Azotobacter vinelandii MoFe protein (NifD/NifK) complexes from wild type (WT) and NifK extension variants (Q45W, KMQ, and NifK-β), along with anaerobically purified NifH (Fe protein). **(b)** Effect of oxygen exposure on nitrogenase activity of NifK variants. **(c)** Effect of oxygen exposure on MoFe oligomeric state, measured by mass photometry. **(d)** Effect of heat exposure (40 °C) on nitrogenase activity of NifK variants. **(e)** Effect of heat exposure on MoFe oligomeric state. Purified MoFe variants were exposed either to atmospheric oxygen (∼21% O₂) or 40 °C for the indicated durations. To assess activity, purified and stress-treated MoFe variants were combined with purified *A. vinelandii* Fe protein and an ATP regeneration system under an atmosphere of 95% N_2_ and 5% H_2_. Nitrogenase activity was then quantified as a measure of ammonia production. Values represent the mean of three technical replicates, with shaded regions indicating ± standard deviation.

To assess the impact of oxidative and heat stress on oligomeric state, purified MoFe variants subjected to stress were analyzed by mass photometry. Prior to stress exposure, all variants exhibit majority tetrameric assemblies (82–95%), except for the Beta deletion variant, which showed minimal tetramer formation (∼7%) (**Fig. S5**). Considering this, we did not include the Beta variant in future mass photometry experiments. Under stress conditions, distinct differences in tetramer stability are observed among the remaining variants. During oxygen exposure, WT (15.5 ± 3.0% loss per hour) and Q45W (11.3 ± 1.4% loss per hour) display similar rates of oligomeric decay, whereas KMQ (36.7 ± 4.0% loss per hour) decays more rapidly. Under heat stress, KMQ again exhibits the fastest decay (56.7 ± 2.5% loss per hour), followed by WT (44.7 ± 2.5%) and Q45W (39.0 ± 2.0%), with Q45W decaying significantly more slowly than WT.

## Discussion

In this study, we demonstrate that structural evolution provides a useful framework for narrowing sequence-function search space and identifying regions that are both functionally important and amenable to engineering. By targeting a lineage-specific structural feature, we constrained our exploration to a small, manageable portion of the nitrogenase sequence–function landscape. Generating variant libraries comprising over 9,000 members within this landscape allowed us to characterize the function of the NifK N-terminus extension, enhance diazotrophic fitness, improve nitrogenase activity, and engineer a more thermostable enzyme.

Our results suggest that the NifK N-terminus extension acts as an interface element that can entrench specific inter-subunit interactions, a concept highlighted in studies of surface-domain evolution (Hochberg et al., 2020). By altering this region, we observed that certain mutations produced epistatic effects, highlighting how these interface domains can constrain evolutionary trajectories while maintaining tolerance to mutation. These findings suggest that such structural features not only effect immediate biochemical function but also influence the evolution of multi-subunit complexes like nitrogenase.

Progressive truncation of the NifK N-terminus extension in *A. vinelandii* revealed that shortening beyond the Beta helix abolishes diazotrophic growth, identifying critical features within the first 35 residues (**Fig. 1d**). Alanine scanning and deep mutational analyses further show that while much of the extension tolerates mutation, a subset of residues are essential for robust growth, especially under elevated temperatures (**Figs. 2c, 3**). Together, these results establish that the NifK extension is not a dispensable appendage, but a critical player in nitrogenase function.

At the same time, the extension is well suited for engineering. Most substitutions are neutral or beneficial, generating a flexible sequence-function landscape. Constrained residues often interface directly with NifD, preserving essential interactions, while surrounding positions can be modified to improve performance. Notably, variant KMQ demonstrates that substantial gains in catalytic activity can be obtained by introducing changes without disrupting these core interactions. In this case, catalytic improvement is accompanied by a tradeoff with complex stability, as KMQ degrades the fastest under stress conditions (**Fig. 5**). Many beneficial variants exhibit temperature-dependent epistatic effects, where certain mutations enhance function under one condition but reduce it under another (**Figs 3e, 4e**). Together, these findings highlight the NifK extension as a critical yet adaptable module, balancing essential inter-subunit interactions with the flexibility needed for engineering improved nitrogenase variants.

Multiple lines of evidence indicate that the primary function of the NifK extension is to stabilize the MoFe complex by reinforcing the NifD-NifK interface. Structural analysis (PDBePISA) suggests that the extension contributes disproportionately to the interface (see Supplementary Table), and truncation progressively decreases predicted binding energy, coinciding with loss of function. Mutations causing detrimental effects are enriched at the interface and are more pronounced at elevated temperatures, consistent with a stabilizing role. For purified MoFe complexes, tetramer formation is severely impaired in the Beta deletion variant but largely maintained in other variants (**Fig. S5**), and mass photometry confirms differences in oligomeric stability under stress with variant Q45W displaying enhanced thermostability. Electrostatic analyses further support co-evolved interaction surfaces. As the extension lengthens, negatively charged regions on NifD progressively redistribute and consolidate at the tip of the N-terminus extension, forming a stable, negatively charged pocket in extant *A. vinelandii* (**Fig. 2b**). Concurrently, the enrichment of positively charged residues within the Beta helix of the extension, including residues predicted to form salt bridges, further supports a co-evolving electrostatic interaction surface (**Fig. 2 b,c**).

Taken together, structural interface analyses, mutational fitness landscapes, enzymatic activity, and oligomeric stability measurements converge to support a stabilizing role for the NifK N-terminus extension, explaining its essentiality and evolutionary conservation. This reveals a structurally critical yet tunable region of nitrogenase that may be leveraged to engineer improved enzymes for biological nitrogen fixation in agriculture.

## Methods and Materials

### Sequence, structure and interface analysis

All ancestral and truncation variant structures were predicted using AlphaFold3 (Abramson et al., 2024), and all NifK lengthening variant structures were predicted using Boltz-2 (Passaro et al., 2025). Structural analyses and electrostatic surface maps were generated in PyMOL. Predicted binding energies between NifD and NifK were obtained using the PRODIGY webserver (Xue et al., 2016). Values for extant *A. vinelandii* NifD and NifK were calculated from the high-resolution crystal structure (PDB: 3U7Q), whereas ancestral and variant values were calculated from their corresponding predicted structures. Interfacing residues between *A. vinelandii* NifD and NifK were identified using the PDBePISA tool applied to PDB 3U7Q.

Extant NifK protein sequences were collected and aligned using previously described methods (Sobol et al., 2026).

### Azotobacter vinelandii transformation

Transformation of *A. vinelandii* was performed following Page and Tigerstrom (1979) and Dos Santos (2019) with substantial modifications. Competent cells were prepared by inoculating competency medium to an initial OD_600_ of 0.01 and growing cultures at 30 °C, shaking at 250 rpm until reaching an OD_600_ of 0.2-0.4. At the desired density, cultures were adjusted to 7% (vol/vol) DMSO, aliquoted into 50 µL volumes, and stored at −80 °C.

For transformation, competent cells were thawed at room temperature and mixed with 500 ng of linearized DNA and 50 µL of 1X MOPS buffer per 50 µL of competent cells. The mixture was incubated at 30 °C for 20 min, after which 50 µL of Burk’s medium (Burk & Lineweaver, 1930) supplemented with nitrogen (BN) were added and cells were allowed to recover for 2 hours at 30 °C while shaking at 250 rpm. Recovered cultures were plated onto a relevant media source and incubated at 37 °C for 2-3 days to obtain isolated colonies.

Newly generated strains were maintained by inoculating 2 mL BN cultures from single colonies and incubating them for 24 hours at 30 °C, shaking at 250 rpm. For long-term storage, 930 µL of each culture were mixed with 70 µL of DMSO and frozen at -80 °C.

### Diazotrophic growth assays

To assess the diazotrophic growth of individually generated NifK variants, each strain was inoculated into 2 mL BN seed cultures in and grown at 30 °C, 250 rpm for 24 hours. Seed cultures were then used to inoculate Burk’s media without nitrogen (B) to an initial OD_600_ of 0.01, followed by incubation at 30 °C. For truncation variants, growth was monitored in triplicate in 50 mL of B media contained within 125 mL baffled flasks shaking at 250 rpm. OD_600_ measurements were taken manually using cuvettes and a UV-vis spectrophotometer. For the NifK point mutants and lengthening variants, growth was measured in quadruplicate in 535 uL B media within 48-well plates using a BMG Labtech plate reader shaking at 300 rpm.

Growth rates were calculated from the exponential portion of each growth curve using equation 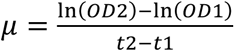, where OD_1_ and OD_2_ are optical densities measured at times t_1_ and t_2_, respectively.

### NifK point-mutant library generation

The NifK point-mutant library was generated using an oligo-pool-based approach following Freschlin et al. (2025) with modifications. Synthetic ssDNA corresponding to each desired variant was ordered as an oligo pool (Twist Bioscience). These oligos were designed to include BsaI sites flanked by primer binding regions for subsequent amplification and Golden Gate assembly. The pool was amplified into dsDNA via PCR using KAPA HiFi HotStart ReadyMix (Roche) with standard cycling conditions and 20 amplification cycles to minimize PCR bias. PCR products were purified using the Zymo DNA Clean and Concentrator kit.

Golden Gate reactions were assembled using an 18:1 molar ratio of amplified oligo insert to insertion vector, with 75 ng of vector backbone per reaction. Each reaction contained 15 U BsaI-HFv2 (NEB) in 2X T4 DNA Ligase Buffer (NEB) and was incubated at 37 °C for 2 hours for digestion. Reactions were then supplemented with 1000 U T4 DNA Ligase (NEB) and incubated for an additional 18 hours at 37 °C for ligation. Enzymes were then heat inactivated at 65 °C for 15 minutes.

Five microliters of the assembly reaction were transformed into chemically competent *E. coli* DH5α cells via standard heat-shock transformation. Transformation was performed at a scale sufficient to obtain a 20X coverage of the library. Following recovery, transformed cells were pooled directly into 50 mL of LB supplemented with carbenicillin (100 µg/mL) and kanamycin (50 µg/mL) and grown for 16 hours at 37 °C, shaking at 250 rpm. Plasmids were isolated from the resulting culture, linearized via digestion with ScaI-HF (NEB) and transformed into *A. vinelandii* for genomic integration at a scale to achieve ∼35X coverage of the library.

### NifK extension lengthening library generation

The NifK extension lengthening library was generated using a similar approach to that used for the point-mutant library, with key modifications to introduce N-terminus extensions. Synthetic ssDNA oligos were ordered as primers from IDT containing degenerate codons and flanking BsaI sites. These primers were used to amplify the NifK transformation vector, thereby appending three randomized codons between positions Q3 and Q4 near the N-terminus of the NifK extension. PCR amplification was performed using Q5 High-Fidelity DNA Polymerase (NEB) under standard cycling conditions with 20 cycles to minimize PCR bias. Amplified products were purified and assembled via Golden Gate cloning using 100 ng of purified PCR product, following the same digestion and ligation conditions described above.

Homologous recombination fragments were generated by PCR amplification directly from the Golden Gate assembly mixture using 1 ng of assembly product as template and Q5 High-Fidelity DNA Polymerase (NEB) with standard cycling conditions and 20 amplification cycles. The resulting fragments were then transformed into *A. vinelandii* for genomic integration at a scale to obtain ∼12X coverage of the library.

### Pooled diazotrophic growth competitions

To assess the diazotrophic fitness of variants in the NifK point-mutant and extension lengthening libraries, pooled growth competition assays were performed. *A. vinelandii* recovery cultures were combined and inoculated into 50 mL of BN medium supplemented with kanamycin (0.6 µg/mL) in triplicate. Cultures were grown at 30 °C with shaking at 250 rpm until reaching an OD₆₀₀ of 2.0. From each culture, 2 mL input samples were collected, harvested by centrifugation at 16,000 × g for 1 min, and stored at −80 °C.

Input cultures were then used to inoculate 50 mL B medium to an initial OD₆₀₀ of 0.01 in triplicate and grown at 20, 30, or 40 °C with shaking at 250 rpm. Cultures were allowed to grow for approximately seven doublings (final OD₆₀₀ ≈ 1.3). From each culture, 2 mL output samples were collected, harvested by centrifugation at 16,000 × g for 1 min, and stored at −80 °C.

Genomic DNA was extracted from frozen cell pellets using the Qiagen DNeasy UltraClean Microbial Kit. NifK extension regions were amplified from 10 ng of genomic DNA per sample using primers containing adapters for index PCR and Illumina sequencing. PCR amplification was performed using Q5 High-Fidelity DNA Polymerase (NEB) under standard cycling conditions with 25 cycles. Amplicons were purified using the Invitrogen PureLink PCR Purification Kit and submitted to the University of Wisconsin sequencing core for index PCR and Illumina sequencing.

### Relative fitness calculations

Raw Illumina sequencing reads were paired, merged, and trimmed using Geneious. Reads corresponding to each variant were counted for every sample, and variant abundance was calculated as the fraction of total reads within each sample. Variants that did not have 100 or more reads from the input population were not included in the analysis. Abundance values were then normalized to wild-type (WT). Relative fitness for each variant was calculated as the log₂ ratio of output relative abundance to input relative abundance for each sample. Positive relative fitness values indicate variants with higher fitness than WT, whereas negative values indicate reduced fitness relative to WT:

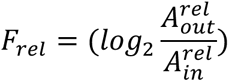

To enable direct comparison between pooled and individual diazotrophic growth assays, growth rates from the individual assays were calculated using a similar formula:

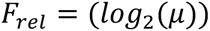

### Individual variant generation

Individual NifK variants were generated by designing primers to either introduce specific amino acid substitutions or insert three additional residues at the N-terminus of the NifK transformation plasmid. Site-directed mutagenesis (SDM) was performed following the NEB Q5 Site-Directed Mutagenesis Kit protocol. Five microliters of each assembly reaction were transformed into chemically competent *E. coli* DH5α cells via standard heat-shock transformation. Following recovery, cells were plated onto LB agar supplemented with carbenicillin (100 µg/mL) and kanamycin (50 µg/mL) and incubated overnight at 37 °C. Individual transformants were grown in 5 mL LB + Kan + Carb cultures overnight at 37 °C, and plasmids were subsequently extracted. Sequence-verified plasmids were then linearized and transformed into *A. vinelandii* for genomic integration.

### NifDK-complex purification

Wild-type *Azotobacter vinelandii* NifDK (WT), extension mutants (Q45W and KMQ), and the NifK-β truncation mutant were purified under strictly anaerobic conditions in a COY anaerobic chamber. The NifD subunit in all constructs were engineered with an N-terminal Strep-tag II, enabling co-purification of the NifDK complex via affinity chromatography. All buffers were rendered anaerobic by equilibration under stirring in the anaerobic chamber for 12 h prior to use, and reducing conditions were maintained by supplementation with sodium dithionite (400 mg/L) where specified.

For WT and NifK extension mutants, cells were grown in BN medium at 30 °C with shaking for 24 h (OD₆₀₀ ≈ 2.5), followed by subculturing into B medium at an initial OD₆₀₀_nm_ of 0.005 and growth at 30 °C, 250 rpm to an OD₆₀₀_nm_ of 2.0. Cells were harvested by centrifugation at 7,500 rpm for 15 min at 4 °C using a JLA 8.1 rotor in a Beckman Coulter Avanti JXN-26 centrifuge. For the NifK-β truncation mutant, which does not grow in nitrogen-free medium, cells were initially cultured in 6 L BN medium to OD₆₀₀_nm_ ≈ 1.0, harvested, and resuspended in 6 L nitrogen-free Bruke’s medium, followed by incubation at 30 °C with shaking for 4 h to induce nitrogenase expression prior to harvest.

Cell pellets were transferred into the anaerobic chamber, resuspended in anaerobic glycerol buffer [4.0 M glycerol, 50 mM Tris-HCl (pH 7.9)], and re-pelleted at 8,000 rpm for 25 min at 4 °C (JL-20 rotor). The pellets were resuspended in anaerobic extraction buffer at five times the pellet volume (e.g., 20 mL buffer per 4 g of pellet) containing 50 mM Tris-HCl (pH 7.9), 0.2 mM phenylmethanesulfonyl fluoride (PMSF), 2 µM pepstatin, 6 U/mL DNase, and 400 mg/L sodium dithionite, followed by cell lysis via osmotic shock. Lysates were clarified by centrifugation at 18,900 rpm for 50 min at 4 °C (JLA-20 rotor), yielding a brown supernatant that was loaded onto a Strep-Tactin®XT 4flow high-capacity resin (IBA Life Sciences, Catalog number: 2-5030-010) pre-equilibrated with 15 column volumes (CV) of anaerobic equilibration buffer [50 mM Tris-HCl (pH 7.9)] in the COY anaerobic chamber.

The column was washed with 10 CV of equilibration buffer (until the flow through from the column turns clear), and the resin bound NifDK complex was eluted with 19 mL elution buffer [50 mM Tris-HCl (pH 7.9), 150 mM NaCl, 50 mM biotin]. Eluted fractions were pooled and concentrated using Amicon Ultra-15 centrifugal filters (Ultracel-3K), and the purified protein was collected in sealed serum vials within the COY anaerobic chamber and stored at -80 °C. Protein concentrations were determined using a BCA assay according to the manufacturer’s instructions.

### NifH-purification

NifH was engineered with an N-terminal Strep-tag II, analogous to NifD, and purified using a protocol similar to that employed for the NifDK complex described above. Briefly, NifH was affinity-purified using Strep-Tactin®XT 4flow high-capacity resin, and elution fractions were concentrated to 3 mL. The concentrated sample was subsequently loaded onto a HiLoad 16/60 Superdex 200 size-exclusion column pre-equilibrated with Buffer A [50 mM Tris-HCl (pH 7.9), 500 mM NaCl, 400mg/L Sodium Dithionite]. Eluted fractions corresponding to NifH were pooled and concentrated using Amicon Ultra-15 centrifugal filters (Ultracel-3K). The purified protein was aliquoted into 50 µL volumes in sealed serum vials and stored at −80 °C.

### Stress Treatment of Purified MoFe Protein

Purified MoFe protein was diluted to the desired concentration in elution buffer lacking sodium dithionite and transferred into Corning® internal-thread cryogenic vials equipped with silicone washers. All sample preparation and handling were performed in a Coy anaerobic chamber under an atmosphere of 95% N₂ and 5% H₂.

For oxygen stress assays, sealed vials were removed from the anaerobic chamber, opened, and exposed to ambient atmosphere (∼21% O₂) for the indicated durations. Following exposure, samples were returned to the anaerobic chamber and resealed prior to downstream analysis.

For heat stress assays, sealed vials were removed from the anaerobic chamber while maintaining anaerobic conditions and incubated at 40 °C for the indicated durations. After incubation, samples were immediately placed on ice.

### In-vitro activity assays

Nitrogenase activity assays were performed as described in Harris et al. (2024) with the following modifications. Reactions contained 0.1 mg/mL MoFe protein (NifD₂K₂) and 0.5 mg/mL Fe protein (NifH₂) in a total volume of 100 µL nucleotide regeneration buffer.

Instead of 9.4 mL vials, assays were conducted in clear, round-bottom 96-well plates. Plates were sealed within a BD GasPak™ EZ Pouch System prior to removal from the anaerobic chamber to maintain an oxygen-free environment. Reactions were incubated at 30 °C with shaking at 250 rpm for 8 minutes and subsequently quenched by the addition of 30 µL of 400 µM EDTA (pH 8.0).

Ammonia production was quantified based on the o-phthalaldehyde (OP) fluorescence assay as described by Corbin (1984) and Harris et al. (2024), with modifications. Briefly, 2.5 µL of the quenched reaction was added to 100 µL of OP assay buffer in a black, flat-bottom, optically clear 96-well plate and incubated at room temperature for 30 minutes in the dark. Fluorescence was measured using a BMG Labtech CLARIOstar Plus plate reader with an excitation wavelength of 410 nm and an emission wavelength of 472 nm.

### Mass photometry

Mass photometry measurements were conducted using a TwoMP mass photometer (Refeyn). Data was collected and analyzed using AcquireMP (Refeyn) and DiscoverMP (Refeyn) respectively. Mass photometry samples were prepared according to methods described by Schmidt et al., 2023.

## Acknowledgements

This work was supported by the National Aeronautics and Space Administration (NASA), the Hypothesis Fund and the W. M. Keck Foundation. The authors thank all Kacar lab members for their support and feedback, especially Bruno Cuevas Zuviría, Katsumi Hagino, Holly Rucker, Evrim Fer, Kaustubh Amritkar, and Trenton Laack. We also thank members of the Seefeldt lab, specifically Derek F. Harris, for providing us with nitrogenase purification and ammonia production assay protocols as well as samples of purified protein for troubleshooting our ammonia production assay; the University of Wisconsin-Madison Biotechnology Center Animal Models Core and Advanced Genome Editing Laboratory (RRID:SCR_021070) for their contributions in generating genome edited models.

Mass photometry data were obtained at the University of Wisconsin – Madison Biophysics Instrumentation Facility, which was established with support from the University of Wisconsin – Madison and grants BIR-9512577 (NSF) and S10 RR13790 (NIH).

## Notes

### Competing Interest Statement

The authors have declared no competing interest.

### Summary of Updates

Updated abstract to remove typos. No other changes.

