## Supplemental Information for "Evolution-guided engineering of an ancient nitrogenase interface enhances enzyme activity and stability"

*Structural analysis of NifK extension aromatic-aromatic interactions*

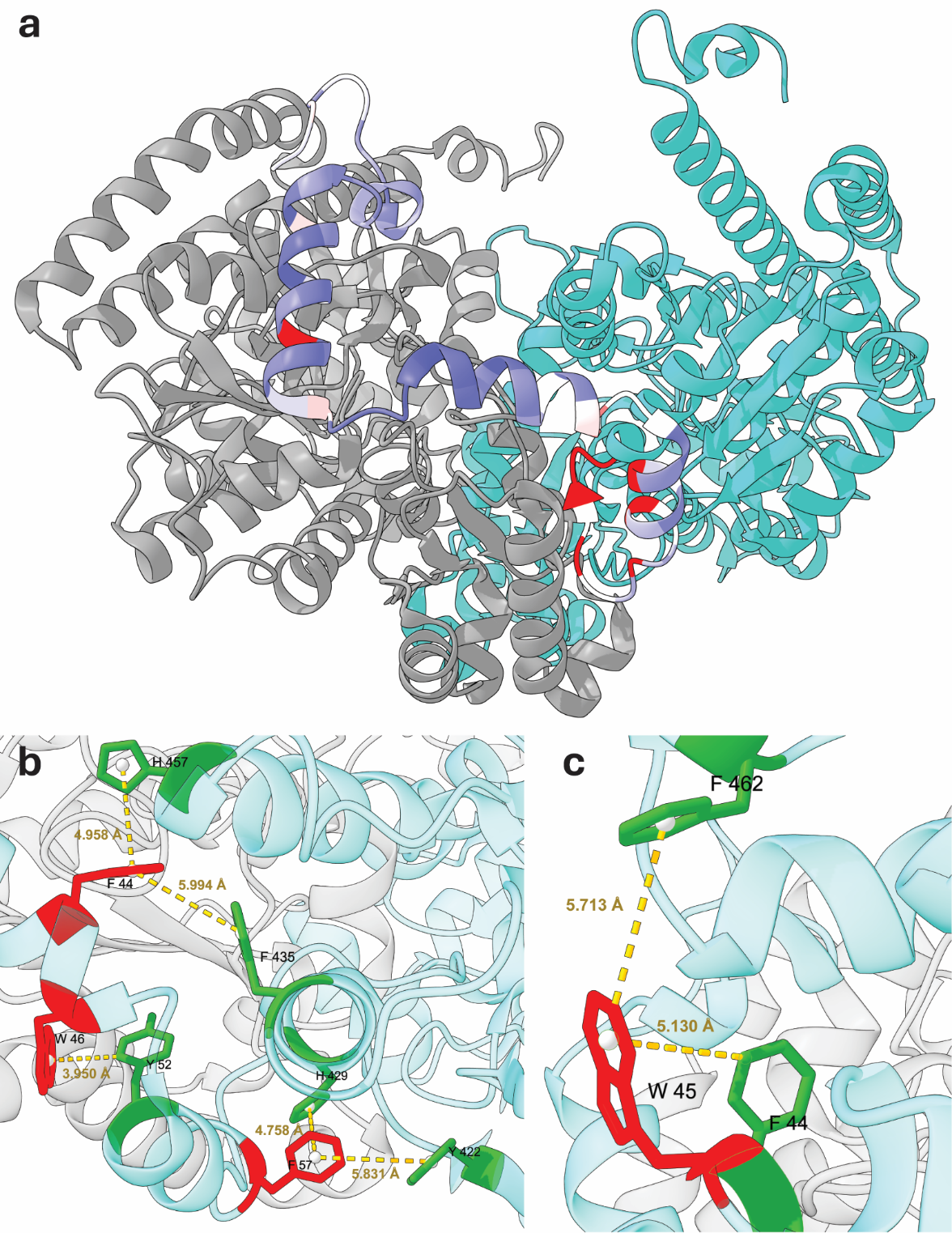

**Fig. S1: Structural analysis of the NifK extension and key residues.** (a) Structure of *A. vinelandii* NifD and NifK (PDB: 3U7Q), with extension residue positions colored by average relative fitness values at 40 °C. (b) Temperature sensitive aromatic region of NifK extension. Residues contributing to temperature sensitivity are shown in red, while potential interacting residues are highlighted in green. (c) Structure of point mutant Q45W. Mutant residue is shown in red, while new potential interactions are highlighted in green.

*Temperature dependence of NifK extension point mutants*

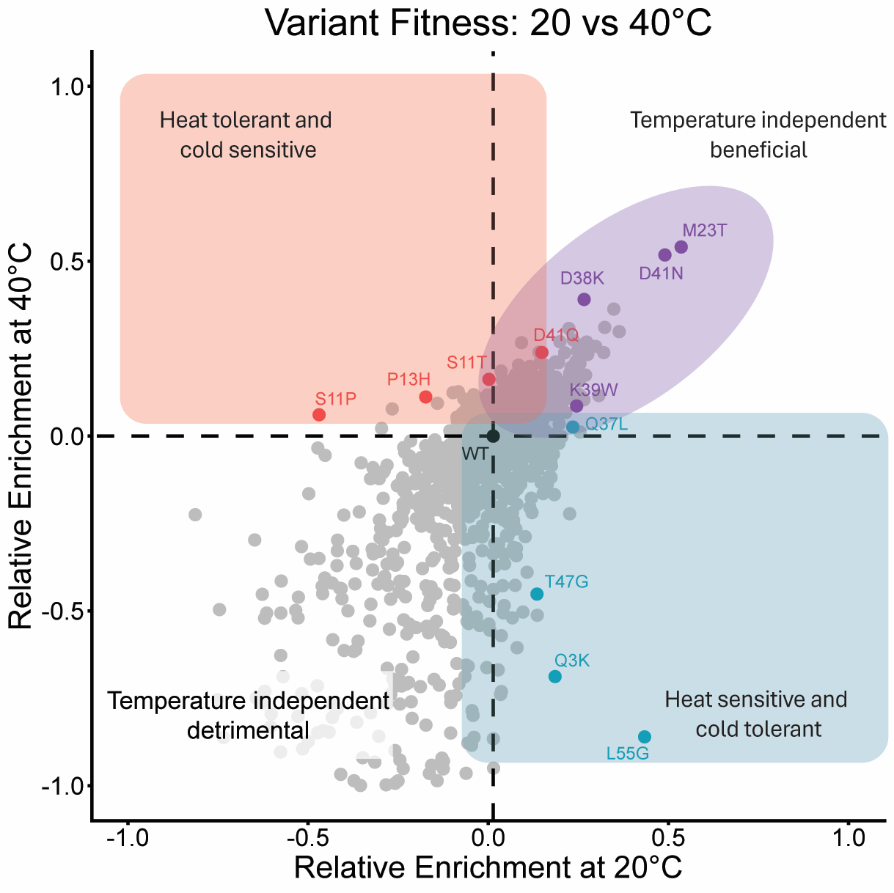

**Fig. S2: Temperature-dependent fitness effects of NifK extension point mutants.** Comparison of relative fitness scores at 20 and 40 °C. The x-axis represents mean fitness at 20 °C and y-axis represents mean fitness at 40 °C. Selected variants are labeled by mutation and colored according to temperature tolerance.

*Combining beneficial NifK mutations*

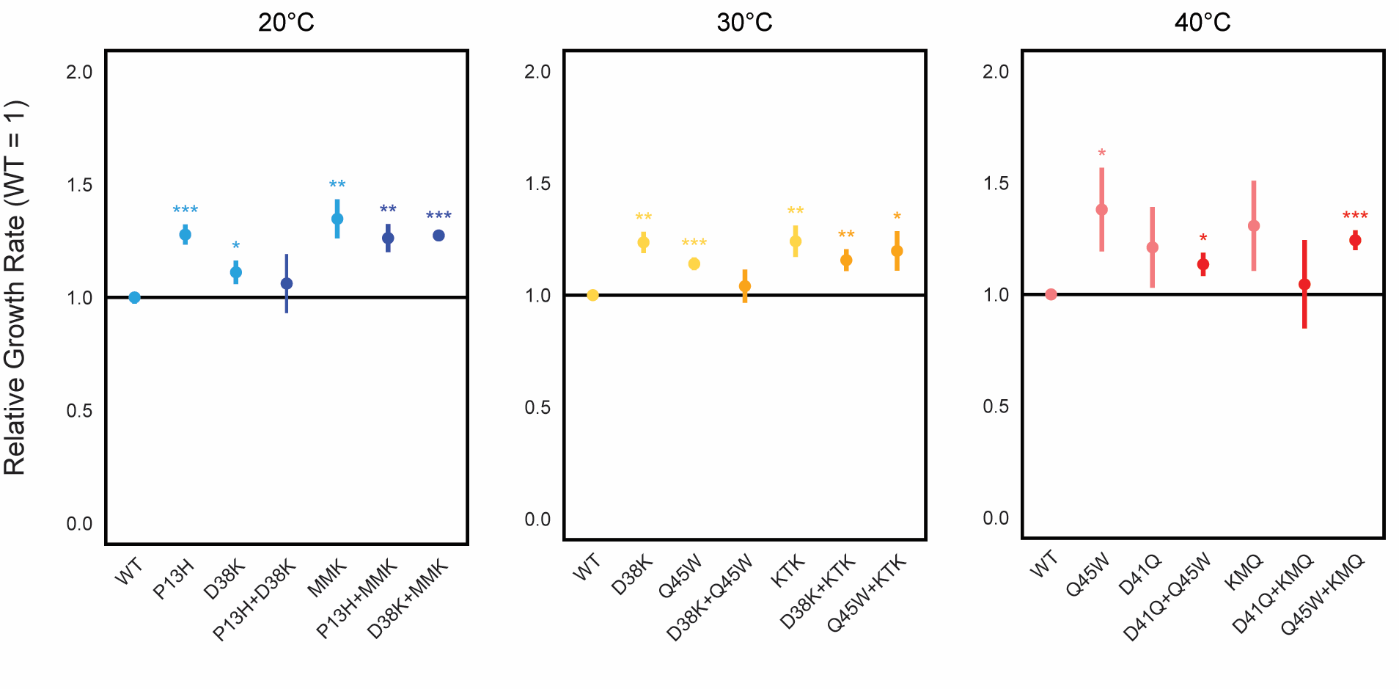

**Fig. S3: Fitness of double mutants is not improved over single mutants.** Comparison of growth rate relative to WT for individually tested single and double mutants at 20, 30 and 40 °C. Values plotted as the mean of four biological replicates. Error bars indicated as the standard deviation of four biological replicates. Significance from WT values displayed as asterisks (* p < 0.05, ** p < 0.01, *** p < 0.001).

*WT vs. KMQ ammonia production assay reproducibility*

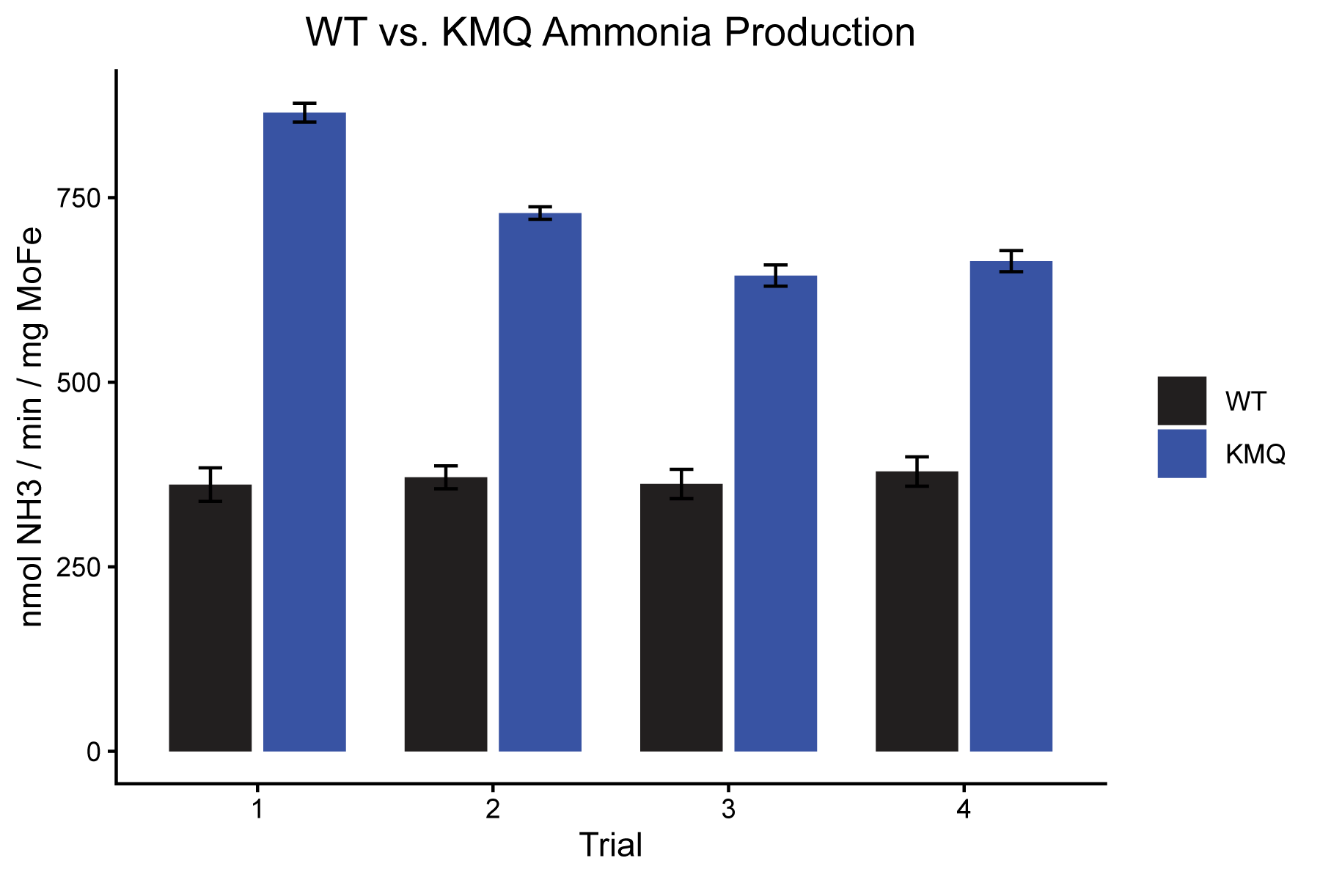

**Fig. S4: Reproducibility of purified KMQ activity.** X-axis represents nitrogenase activity quantified as nmol of ammona produced per minute per milligram of MoFe protein. Y-axis represents data from independent trials.

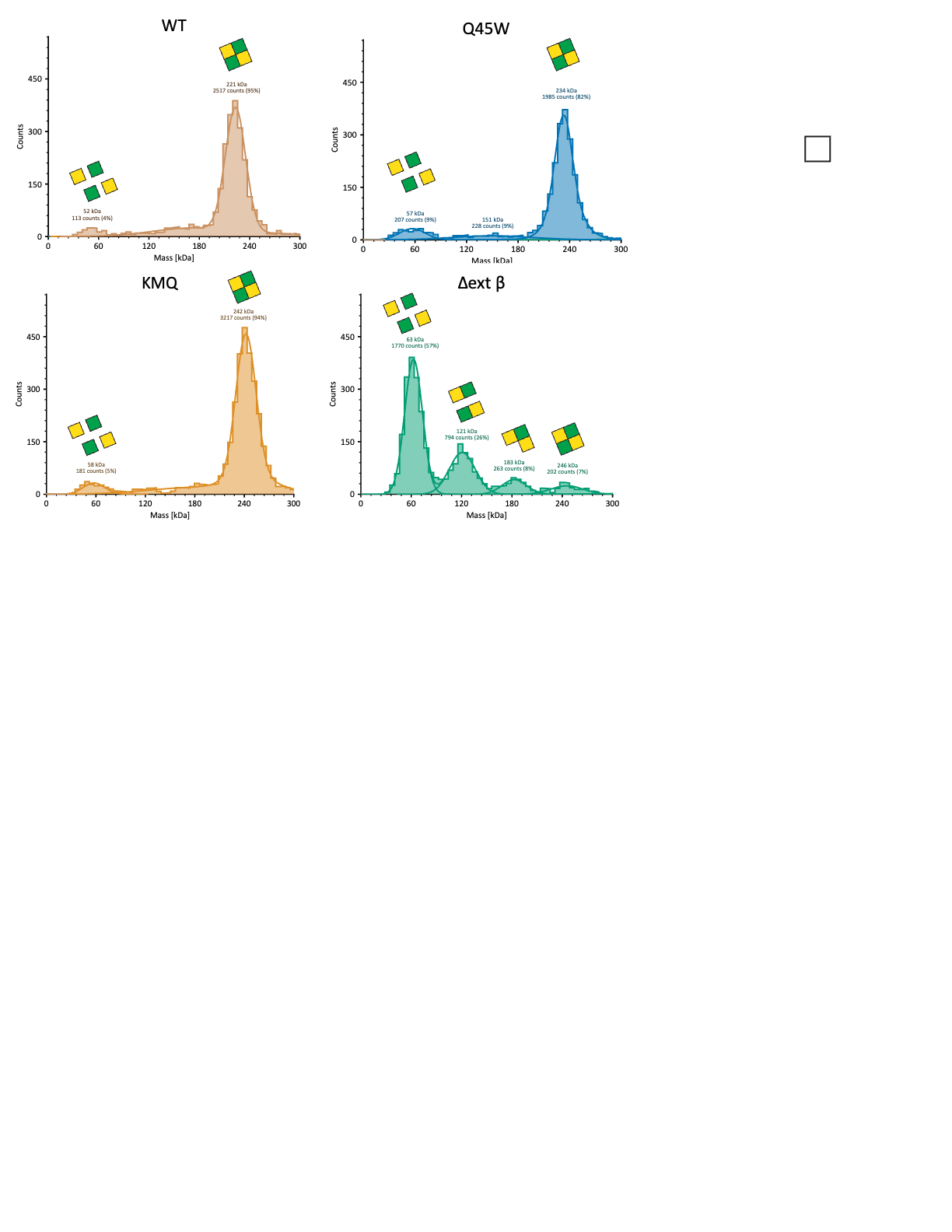
*Oligomeric state of purified NifK variants*

**Fig. S5: Oligomeric state of purified NifK variants.** Mass photometry data of purified MoFe variants. X-axis represents approximate mass of analyze molecules. Y-axis represents the total number of molecular counts during the measurement period. Oligomeric state visualized over respective size peaks. Mass, total count and relative percentage of counts labeled over each peak.

| Variant | Rel. Fitness 20 °C  (DMS / Individual) | Rel. Fitness 30 °C  (DMS / Individual) | Rel. Fitness 40 °C  (DMS / Individual) |
| --- | --- | --- | --- |
| Q3K | 0.17 / 0.18 | -0.03 / 0.44 | -0.69 / -0.09 |
| S11T | -0.01 / 0.26 | 0.07 / 0.24 | 0.16 / 0.02 |
| S11P | -0.48 / 0.11 | 0.05 / 0.63 | 0.06 / -0.10 |
| P13H | -0.19 / 0.44 | 0.04 / 0.54 | 0.11 / 0.26 |
| K21A | -0.06 / -0.20 | 0.33 / 0.32 | -0.08 / 0.16 |
| M23T | 0.52 / 0.19 | 0.22 / 0.58 | 0.54 / 0.32 |
| Q37L | 0.22 / -0.19 | 0.13 / 0.11 | 0.03 / -0.01 |
| D38K | 0.25 / 0.36 | 0.36 / 0.88 | 0.39 / 0.42 |
| K39W | 0.23 / -0.13 | 0.19 / -0.09 | 0.09 / -1.17 |
| D41N | 0.48 / 0.18 | 0.52 / 0.27 | 0.52 / 0.30 |
| D41Q | 0.14 / 0.21 | 0.28 / 0.49 | 0.24 / 0.77 |
| Q45E | 0.05 / -0.09 | 0.32 / 0.62 | 0.15 / 0.80 |
| Q45W | 0.17 / -0.30 | 0.41 / 0.92 | 0.11 / 0.87 |
| T47G | 0.12 / -0.15 | 0.12 / 0.25 | -0.45 / -1.63 |
| L55G | 0.42 / 0.18 | -0.12 / 0.35 | -0.86 / -0.02 |
| NWM | 1.62 / 0.14 | 1.79 / 0.14 | 1.17 / 0.06 |
| KTK | 1.61 / 0.38 | 1.39 / 0.77 | 1.06 / 0.21 |
| NQS | 1.52 / 0.37 | 1.14 / 0.72 | 0.94 / 0.09 |
| PYM | 1.01 / -0.23 | 1.45 / 0.16 | 0.97 / 0.01 |
| MMK | 1.44 / 0.50 | 1.42 / 0.27 | 0.57 / -2.00 |
| NCK | 1.18 / -0.08 | 1.29 / 0.04 | 0.63 / -0.13 |
| KFS | 1.67 / 0.39 | 1.33 / 0.32 | 1.43 / 0.41 |
| KLD | 1.14 / 0.32 | 1.20 / 0.32 | 1.25 / 0.08 |
| KMQ | 1.23 / -0.22 | 0.95 / 0.33 | 1.15 / 0.54 |

**Table S1: Relative fitness values of individually tested mutants.** DMS fitness values represented as mean of three biological replicates. Individual fitness values represented as mean of four biological replicates.

|  | Position #1 | | | Position #2 | | | Position #3 | | |
| --- | --- | --- | --- | --- | --- | --- | --- | --- | --- |
| Amino Acid | 20 °C | 30 °C | 40 °C | 20 °C | 30 °C | 40 °C | 20 °C | 30 °C | 40 °C |
| A | 0.30 | 0.28 | 0.26 | 0.41 | 0.42 | 0.43 | 0.38 | 0.39 | 0.40 |
| C | 0.41 | 0.43 | 0.31 | 0.44 | 0.43 | 0.35 | 0.43 | 0.44 | 0.35 |
| D | 0.53 | 0.47 | 0.43 | 0.51 | 0.48 | 0.41 | 0.51 | 0.53 | 0.43 |
| E | 0.35 | 0.43 | 0.37 | 0.48 | 0.47 | 0.39 | 0.47 | 0.46 | 0.35 |
| F | 0.51 | 0.52 | 0.36 | 0.51 | 0.52 | 0.37 | 0.50 | 0.52 | 0.37 |
| G | 0.20 | 0.19 | 0.20 | 0.38 | 0.41 | 0.36 | 0.34 | 0.33 | 0.27 |
| H | 0.48 | 0.48 | 0.42 | 0.45 | 0.49 | 0.40 | 0.43 | 0.44 | 0.37 |
| I | 0.45 | 0.52 | 0.38 | 0.49 | 0.52 | 0.41 | 0.47 | 0.49 | 0.43 |
| K | 0.65 | 0.63 | 0.48 | 0.59 | 0.62 | 0.47 | 0.63 | 0.59 | 0.46 |
| L | 0.36 | 0.41 | 0.30 | 0.39 | 0.40 | 0.31 | 0.38 | 0.39 | 0.31 |
| M | 0.45 | 0.50 | 0.37 | 0.43 | 0.45 | 0.34 | 0.49 | 0.52 | 0.39 |
| N | 0.58 | 0.61 | 0.49 | 0.53 | 0.54 | 0.42 | 0.54 | 0.59 | 0.49 |
| P | 0.46 | 0.45 | 0.39 | 0.42 | 0.42 | 0.35 | 0.45 | 0.46 | 0.40 |
| Q | 0.49 | 0.47 | 0.38 | 0.49 | 0.48 | 0.37 | 0.50 | 0.51 | 0.37 |
| R | 0.31 | 0.30 | 0.30 | 0.42 | 0.41 | 0.32 | 0.39 | 0.38 | 0.28 |
| S | 0.44 | 0.45 | 0.36 | 0.43 | 0.45 | 0.33 | 0.45 | 0.47 | 0.35 |
| T | 0.44 | 0.46 | 0.38 | 0.47 | 0.46 | 0.39 | 0.45 | 0.47 | 0.36 |
| V | 0.32 | 0.34 | 0.25 | 0.38 | 0.44 | 0.32 | 0.37 | 0.38 | 0.30 |
| W | 0.41 | 0.42 | 0.37 | 0.41 | 0.45 | 0.38 | 0.37 | 0.40 | 0.31 |
| Y | 0.47 | 0.48 | 0.36 | 0.49 | 0.51 | 0.41 | 0.48 | 0.52 | 0.41 |

**Table S2: Fitness impact of amino acids in beneficial extended NifK variants.** Fitness values of all beneifical variants split into individual positions and averaged by amino acid identity.

|  | Position #1 | | | Position #2 | | | Position #3 | | |
| --- | --- | --- | --- | --- | --- | --- | --- | --- | --- |
| Amino Acid | 20 °C | 30 °C | 40 °C | 20 °C | 30 °C | 40 °C | 20 °C | 30 °C | 40 °C |
| A | 1.30 | 1.62 | 1.76 | 1.76 | 1.93 | 2.01 | 1.56 | 1.81 | 1.87 |
| C | 0.48 | 0.61 | 0.55 | 0.71 | 0.91 | 0.73 | 0.96 | 1.20 | 0.92 |
| D | 1.62 | 1.72 | 1.64 | 1.17 | 1.32 | 1.22 | 1.10 | 1.27 | 1.10 |
| E | 2.69 | 2.71 | 2.40 | 2.04 | 2.18 | 2.07 | 1.54 | 1.71 | 1.58 |
| F | 0.39 | 0.42 | 0.42 | 0.54 | 0.71 | 0.47 | 0.78 | 0.93 | 0.66 |
| G | 1.97 | 2.35 | 2.34 | 2.32 | 2.52 | 2.45 | 1.86 | 2.14 | 2.04 |
| H | 0.49 | 0.67 | 0.58 | 0.68 | 0.79 | 0.69 | 0.94 | 0.98 | 0.89 |
| I | 0.46 | 0.59 | 0.66 | 0.75 | 0.88 | 0.57 | 0.80 | 0.85 | 0.68 |
| K | 1.12 | 1.11 | 0.91 | 1.17 | 1.22 | 1.01 | 1.19 | 1.27 | 1.11 |
| L | 0.86 | 0.91 | 0.91 | 0.80 | 1.01 | 0.97 | 1.11 | 1.31 | 1.25 |
| M | 0.80 | 0.87 | 0.78 | 1.10 | 1.31 | 1.15 | 0.95 | 1.19 | 1.04 |
| N | 0.40 | 0.43 | 0.31 | 0.79 | 0.95 | 0.62 | 0.75 | 0.79 | 0.63 |
| P | 0.75 | 0.97 | 0.89 | 0.89 | 1.09 | 0.94 | 1.16 | 1.28 | 1.09 |
| Q | 1.17 | 1.43 | 1.28 | 1.04 | 1.30 | 1.41 | 1.22 | 1.33 | 1.25 |
| R | 1.34 | 1.57 | 1.46 | 1.24 | 1.49 | 1.38 | 1.46 | 1.69 | 1.50 |
| S | 0.46 | 0.59 | 0.47 | 0.70 | 0.92 | 0.70 | 0.90 | 1.16 | 1.00 |
| T | 0.69 | 0.80 | 0.71 | 1.04 | 1.19 | 1.02 | 0.98 | 1.14 | 0.93 |
| V | 0.89 | 1.12 | 1.16 | 1.34 | 1.46 | 1.51 | 1.28 | 1.45 | 1.35 |
| W | 2.07 | 2.18 | 2.11 | 1.51 | 1.71 | 1.56 | 1.81 | 1.87 | 1.87 |
| Y | 0.40 | 0.42 | 0.37 | 0.62 | 0.76 | 0.49 | 0.82 | 0.91 | 0.67 |

**Table S3: Fitness impact of amino acids in detrimental extended NifK variants.** Fitness values of all detrimental variants split into individual positions and averaged by amino acid identity.
